# Segregostat: A novel concept to control phenotypic diversification dynamics on the example of Gram-negative bacteria

**DOI:** 10.1101/542704

**Authors:** Hosni Sassi, Thai Minh Nguyen, Samuel Telek, Guillermo Gosset, Alexander Grünberger, Frank Delvigne

## Abstract

Controlling and managing the degree of phenotypic diversification of microbial populations is a challenging task. This task not only requires detailed knowledge regarding diversification mechanisms but also advances technical setups for the real-time analyses and control of population behavior on single-cell level. In this work, setup, design and operation of the so called segregostat is described which, in contrast to a traditional chemostats, allows the control of phenotypic diversification of microbial populations over time. Two exemplary case studies will be discussed, emphasizing the applicability and versatility of the proposed approach. In detail the phenotypic diversification of *Eschericia coli* or *Pseudomonas putida* based on monitoring membrane permeability will be controlled. We show that upon nutrient limitation, cell population tends to diversify into several subpopulations exhibiting distinct phenotypic (non-permeablized and permeablized cells). On-line analysis leads to the determination of the ratio between cells in these two states, which in turn trigger the addition of glucose pulses in order to maintain a pre-defined diversification ratio. These results prove that phenotypic diversification can be controlled by means of defined pulse-frequency modulation within continuously running bioreactor setups. This lays the foundation for systematic studies, not only of phenotypic diversification but also for all processes where dynamics single cell approaches are required, such as synthetic co-culture processes.

## 1. Introduction

Controlling and managing the degree of phenotypic diversification of microbial populations has recently attracted a lot of attention [1][2], due to our expanding knowledge about noise of biological systems [3]. This trend can be notably identified by the emergence of new disciplines, such as cybergenetics [4][5], applying control theory for managing cell-to-cell heterogeneity in gene expression at a very high spatio-temporal resolution. A typical device that is thoroughly used for microbial physiology studies is the chemostat allowing the long-term cultivation of microbial population in defined limiting conditions [6]. The main reason behind the intense use of this cultivation device is because it was recognized as being able to stabilize microbial population in a given physiological state. However, results accumulated in recent years pointed out that this is not true [6], and the impact of phenotypic diversification has not been taken into account so far. Indeed, strain-wise, the chemostat is very dynamic. There could be many competing strains, at low frequencies, and a few at higher frequencies, all constantly changing. There can be many successive sweeps (takeovers) that are non-detectable at the OD or substrate levels. Therefore novel tools need to be established based on combination of single cell technologies, such as flow cytometry (FC), and process engineering approaches.

Nutrient stress is a strong trigger of phenotypic diversification that can be eventually used for controlled the degree of diversification of a bacterial population. Previous studies have shown that such phenotypic diversification mechanisms are governed by a complex set of physiological mechanisms involving noise in gene expression, metabolism and growth [7][8][9], these three mechanisms being highly cross-correlated and being controlled through mutual feedback loops. Ultimately, the superimposition of these three mechanisms lead to population phenotypic heterogeneity that can in turn confer interesting functionalities to the whole population (e.g., bet-hedging, division of labor…) [10]. Most of the works focused on phenotypic diversification of microbial populations have been carried out based on single cell proxies involving either GFP expression (used as a proxy for noise in gene expression) [11][12] or growth rate [13][14]. In the context of this work, we propose to focus on another relevant single cell proxy, i.e. membrane permeabilisation. Indeed, membrane permeability is a fundamental physiological parameter driving the way that microbes respond to environmental cues and, eventually, adapt to stresses [15]. However, like growth rate, membrane permeability is a physiological process involving an intricate set of genes and regulation processes. It is thus of importance to develop advanced single cell technologies for characterizing such physiological parameter. This phenomenon is depicted in figure 2 for *Escherichia coli* BW25113 Δ*ompC* and *Pseudomonas putida* KT2440. Propidium iodide (PI) is the most frequently used fluorescence indicator for cell viability based on assessment of membrane integrity [16]. Ordinarily, PI molecule carries two positive charges, one of which seems open to the surroundings and should prevent its membrane permeation. In a few conditions, such as high ATP turnover and nutrient limitation, an increased membrane potential will amplify the ion-motive force for cations, particularly if they carry two charges as the propidium cation [17]. Consequently, a boosted membrane potential might facilitate diffusion of PI molecules inside perisplasmic space. This mechanism allowed us to clearly observe two subpopulations of cells bacteria staining red fluorescent protein (PI+) and bacteria staining very low levels of PI or not staining PI (PI-) even in exponential phase or a feast-and-famine state in glucose pulsed chemostat [18]. In this case, the fraction of the cells that are taking up propidium ions and exhibit intermediate red fluorescence level are not dead and the heterogeneity in physiological states at single cell level can be characterized [19].

It is interesting to point out that outer membrane (OM) permeabilization leads to similar phenotype for our two Gram-negative model organisms. However, if some underlying physiological mechanisms can be advanced for *E. coli*, this physiological process has not been reported for *P. putida*. The OM of Gram negative bacteria is composed of an asymmetric lipid bilayer, phospholipids (PLs) being located in the inner leaflet of the membrane and lipopolysaccharides (LPS) being located to the outer leaflet. In *E. coli*, the transposition of PLs from inner to outer leaflet is obstructed by the MlaA walls surrounding the defined channel. By contrast, outer leaflet PLs can entry this channel and are removed from the OM via transfer to MlaC with the particularly interaction of protein membrane [20][21]. The combined function of porin OmpC and MlaC in lipid transport in maintaining membrane asymmetry has been observed [20]. Addition to the disruption in OM lipid asymmetry, cells lacking OmpC exhibit OM permeability defects, including increased sensitivity to SDS/EDTA and detergents. The authors also suggest that OM permeability defects in Δ*ompC* cells are not simply a result of disruption in OM lipid asymmetry and OmpC may be important for other processes that influence the integrity and function of the OM. Membrane asymmetry is also known to impact different bilayer properties, including cell shape, surface charge, permeability as well as membrane potential [22]. Hence, removing OmpC causes imbalance in OM lipids components that affect cell morphology, membrane integrity and membrane potential. Such increase in permeability for Δ*ompC* mutant has also been observed in this work (Supporting Informations). However, this mechanism seems to be subjected to high cell-to-cell heterogeneity since two clearly defined subpopulations can be observed upon PI staining (Figure 2B). Porins organization and regulation have been less characterized in the case of *P. putida*. By analogy with *E. coli*, these porins are organized in different clusters involving membrane stabilization, cell structure determination, transport of specific substrates and pore formation [23]. Among them, OprB is the porin that has been more thoroughly investigated and present a high homology with OprB from *P. aeruginosa* and has been suggested to be involved in glucose uptake. In *E. coli*, once glucose enters the periplam, it is internalized and phosphorylated by the phosphoenolpyruvate:sugar phosphotransferase system. Catabolism of the generated glucose-6-phosphate proceeds by the glycolytic pathway [24]. In contrast, *P. putida* has an incomplete glycolytic pathway since it lacks 6-phosphofructokinase [25]. Therefore, it metabolizes glucose via the Entner-Doudoroff pathway, where 6-phosphogluconate is the key intermediate. When glucose enters the periplasm in this organism, it can be imported to the cytoplasm via an ABC transporter and then phosphorylated by glucokinase. Glucose is also substrate of the periplasmic enzymes glucose dehydrogenase and gluconate dehydrogenase, yielding gluconate and 2-ketogluconate, respectively. These two compounds and glucose-6-P are the substrates of three convergent pathways leading to the synthesis of 6-phosphogluconate [26].

Porin organization and regulation, as well as metabolic pathways regulation, are fundamentally different between *E. coli* and *P. putida*. However, these microbes display similar features in terms of phenotypic diversification upon nutrient limitation. Indeed, Pi-staining reveals in both cases two clearly defined subpopulations, i.e. the first one exhibiting no PI uptake and the second one exhibiting partial staining probably due to the localization of PI molecules inside the perisplasm upon OM permeabilization (Figure 2).

The dynamics of membrane permeabilisation will be considered as a model system for dynamic high-throughput single cell analyses. More precisely, population profiling by on-line flow cytometry will be performed in chemostat mode. In a second time, subpopulation ratio will be controlled based on a feedback control loop working based on flow cytometry data in a device we called segregostat. According to cybergenetics principle, i.e. applying control theory to cell physiology, and the constraints of gene regulation network, two main control strategies can be envisioned, i.e. either the use of pulse-frequency modulation (PFM) or pulse-width modulation (PWM) [5]. In the context of this work, the first strategies will be used, i.e. glucose pulses for controlling subpopulation ratio of permeabilized and non-permeabilized subpopulations.

## 2. Material and methods

### Strains and medium composition

The strains used in this study are *E. coli* JW2203-1 Δ*ompC* (obtained from the Keio collection [27]. Genotype:Δ*lacZ*4787(::*rrnB*-3), λ^-^, Δ*ompC*768::kan, rph-1, Δ(*rhaD-rhaB*)568, hsdR514) and *Pseudomonas putida* KT2440 (kindly provided by Prof. Pablo I. Nikel, Denmark Technical University, Lyngby). *E. coli* JW2203-1 Δ*ompC* have been selected based on a prescreening test since it was able to display a higher diversification ratio by comparison with wild type and other porin mutants (see Supporting Informations). All strains are maintained at −80°C in working seeds vials (2 mL) in solution with LB media and with 30% of glycerol. Precultures and cultures have been performed on a defined mineral salt medium containing (in g/L): K_2_HPO_4_ 14.6, NaH_2_PO_4_.2H_2_O 3.6; Na_2_SO_4_ 2; (NH_4_)_2_SO_4_ 2.47, NH_4_Cl 0.5, (NH_4_)_2_-H-citrate 1, glucose 5, thiamine 0.01, antibiotic 0.1. Thiamin is sterilized by filtration (0.2 µm). The medium is supplemented with 3mL/L of trace elements solution, 3mL/L of a FeCl_3_.6H_2_O solution (16.7 g/L), 3mL/L of an EDTA solution (20.1 g/L) and 2mL/L of a MgSO_4_ solution (120 g/L). The trace elements solution contains (in g/L): CoCl_2_.H_2_O 0.74, ZnSO_4_.7H_2_O 0.18, MnSO_4_.H_2_O 0.1, CuSO_4_.5H_2_O, CoSO_4_.7H_2_O. The medium was supplemented with 5 g/L of glucose and antibiotic (kanamycin 25 µg/ml) was added for the cultivation of the *E. coli* JW2203-1 Δ*ompC* strain.

Cultivation in bioreactor with on-line flow cytometry (FC) profiling has been performed from overnight precultures performed in 1L baffled flasks containing 100 ml of culture medium and stirred with 200 rpm at 37°C. Cultures in chemostat and segregostat mode have been performed in lab-scale stirred bioreactor (Biostat B-Twin, Sartorius. Total volume: 2L; working volume: 1L). For the batch phase, the overnight cultures were diluted into 1L of minimal medium at an initial OD_600nm_ of 0.5. The pH was maintained at 6.9 by automatic addition of ammonia or phosphoric acid. The temperature was maintained at 37°C under continuous stirring rate of 800 rpm and aeration rate of 1 VVM. Upon glucose depletion, observed typically after 4 – 6 hours with a sudden increase in dissolved oxygen, the chemostat or segregostat mode is started.

For chemostat cultivations, the medium was continuously fed with the complete minimal medium at a dilution rate of 0.1 h^-1^. For segregostat cultivations, the medium was continuously fed with salt basal minimal medium except the carbon source (glucose) at a dilution rate of 0.1 h^-1^. The pulse of glucose was fed in the culture medium according to the regulation sequence controlled by the online software, as it will be discussed further.

The experimental platform is an improved version of a previous on-line flow cytometry platform [28][12] and actually comprises three modules (Figure 1): (a) a conventional culture device (Biostat B-Twin, Sartorius, 2 L), a physical interface for sampling and dilution comprising peristaltic pumps and a mixing chamber (c) detection device, i.e. an Accuri C6 flow cytometer (BD Accuri, San Jose CA, USA). Either module b or c is operated via custom C++ script.

**Figure 1:**
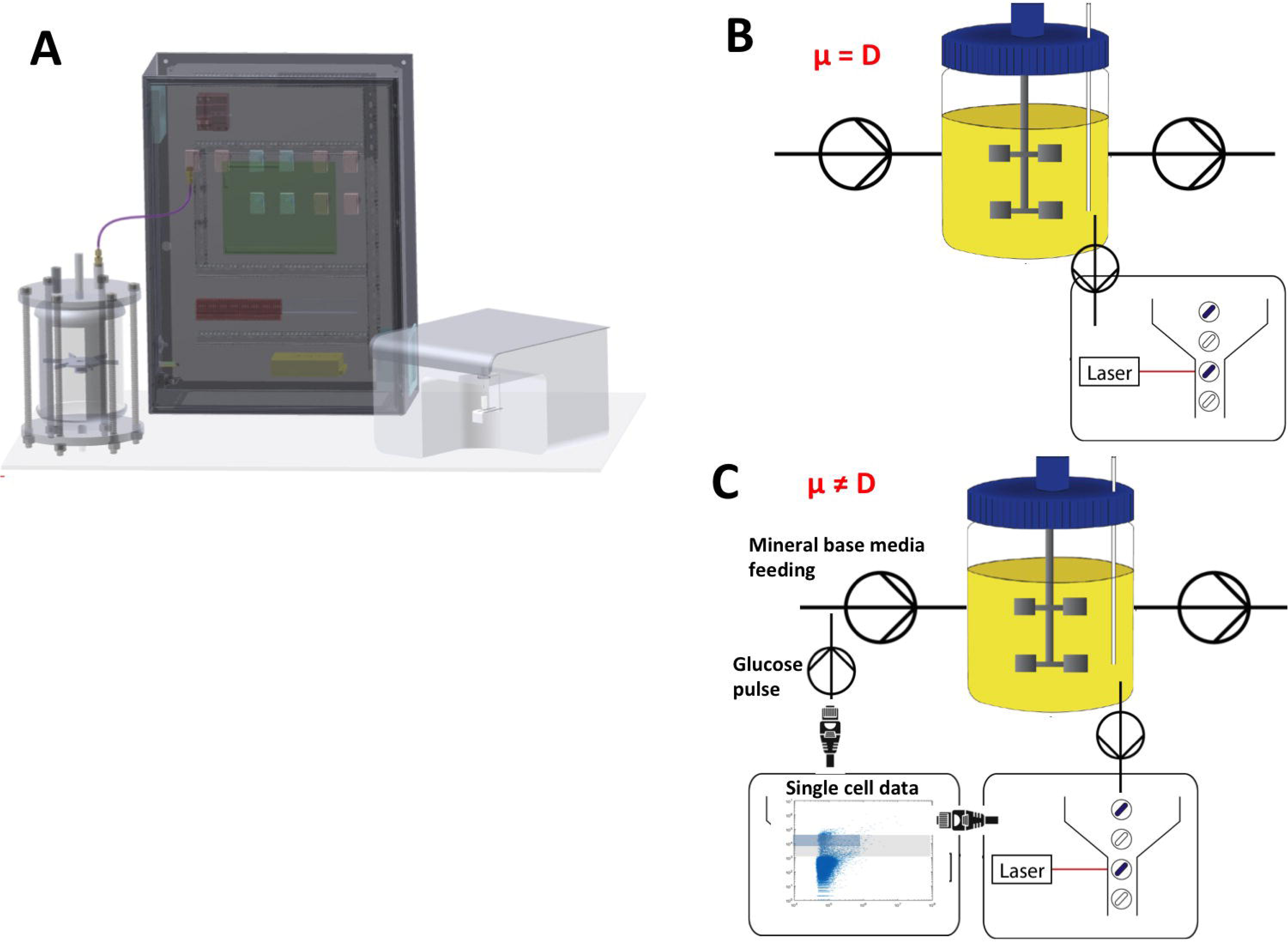
A scheme with the different components for the on-line FC platform. B use of the platform in the chemostat mode. C use of the platform in the segregostat mode. It is important to keep in mind that during segregostat glucose pulses is added based on population diversification and therefore the growth rate (μ) is not necessarily equal to the dilution rate (D).

**Figure 2:**
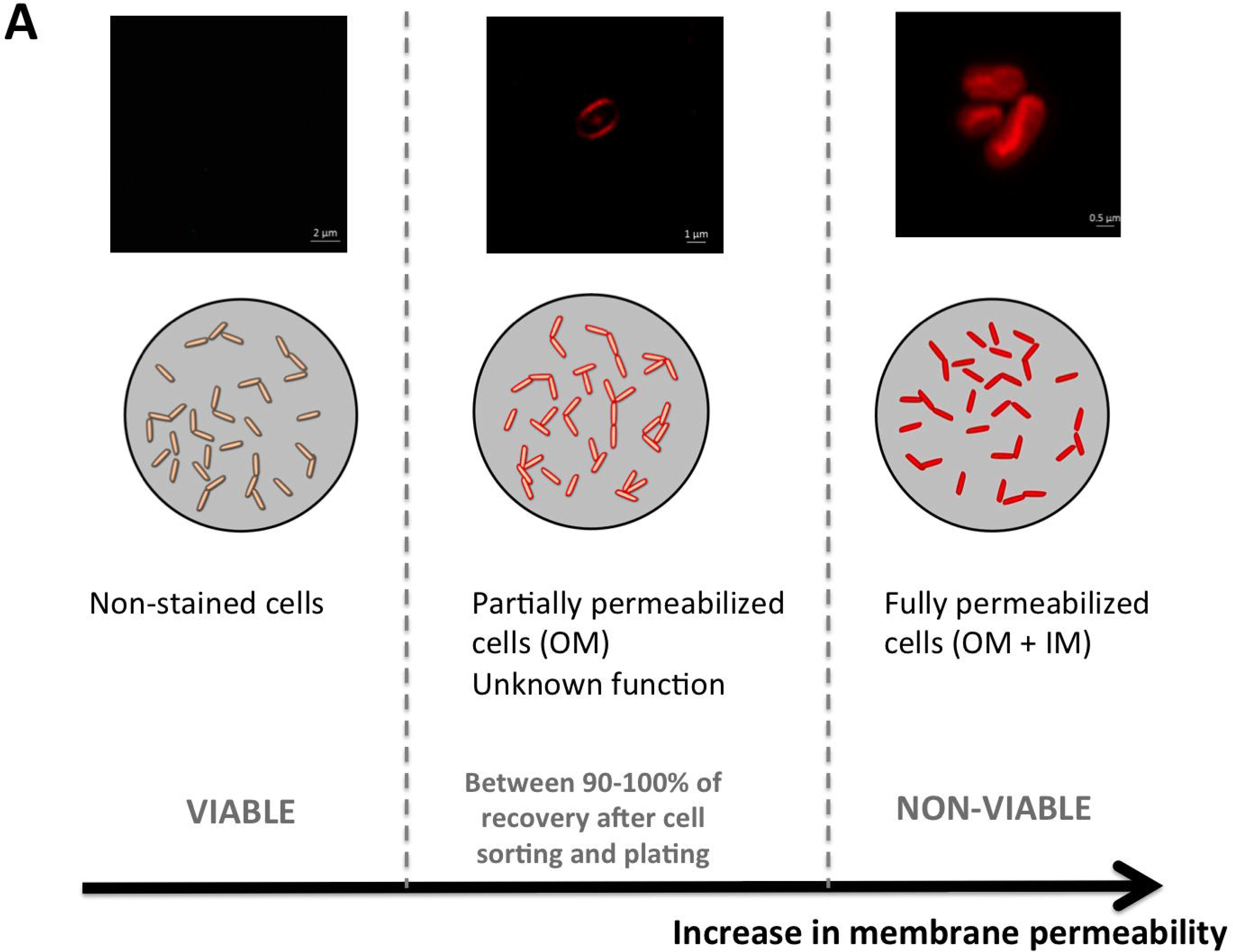
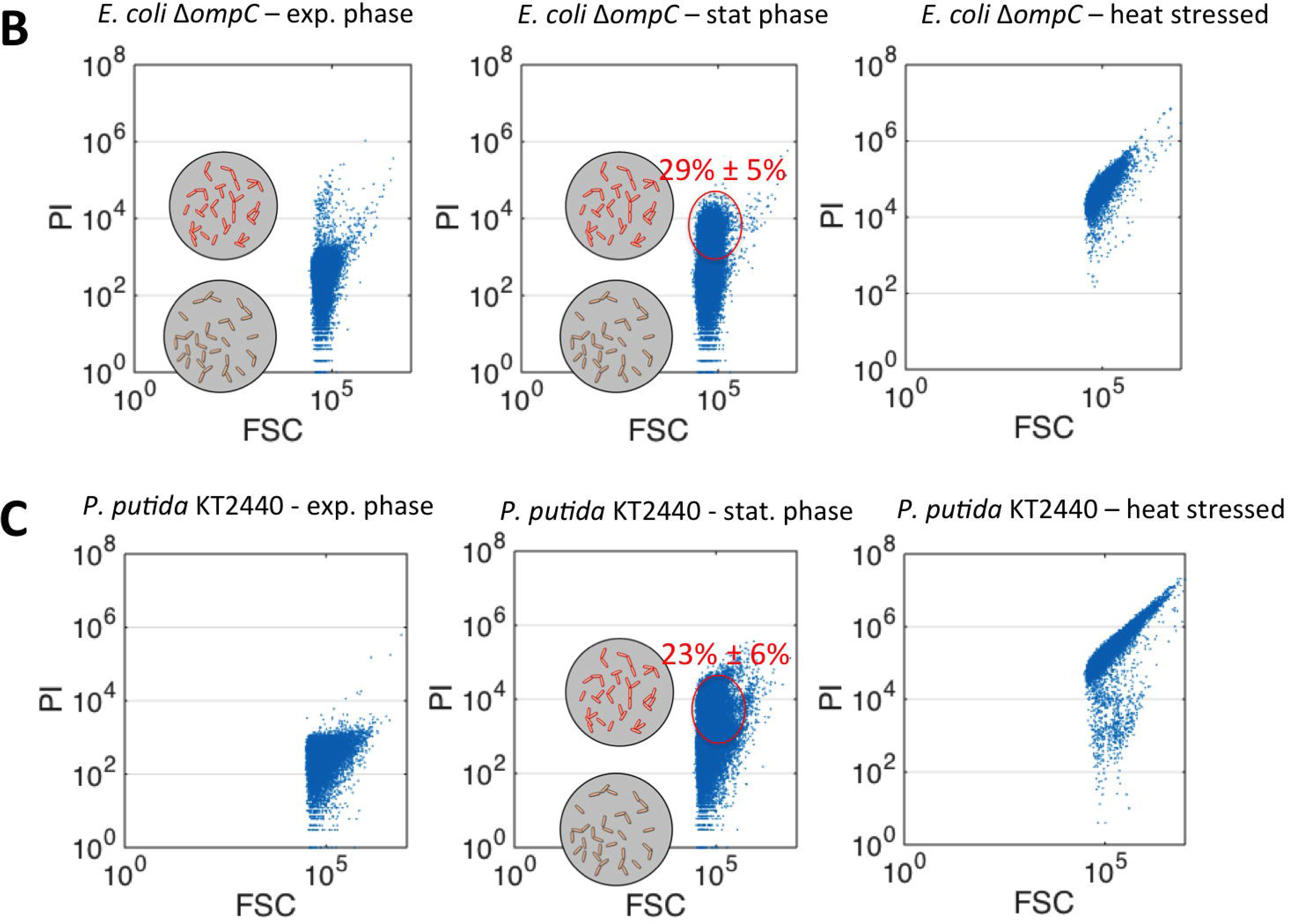
A with the three physiological state according to PI staining (each state is illustrated by images acquired by high resolution microscopy (confocal microscopy with Ayriscan detector). B Flow cytometry experiments (x-axis corresponding to forward scatter or FSC; y-axis corresponding to red fluorescence related to PI staining) with *E. coli* taken at different cultivation stages (cultures made in flasks). C Flow cytometry experiments (x-axis corresponding to forward scatter or FSC; y-axis corresponding to red fluorescence related to PI staining) with *P. putida* taken at different cultivation stages (cultures made in flasks).

In short, the script for processing each sample comprises the following steps: (1) sample acquisition and online staining, (2) online FC analysis, (3) dilution threshold and (4) feedback control loop.

The sample is fed and removed from the mixing chamber based on silicone tubing (internal diameter: 0.5mm; external diameter: 1.6mm, VWR, Belgium) and five peristaltic pumps (400FD/A1 OEM-pump ∼13 rpm and 290 rpm, Watson Marlow). Before and after each experiment, all the connection part (tubing, pumps and mixed chamber) are continuously cleaned with ethanol and rinsed with filtered PBS.

The sample from bioreactor was drawn every 12-min intervals from the bioreactor according to the set of dilution sequences that are controlled via the online software. The latter is working through the following sequence of steps. Firstly, the sampling tube was automatically purged for 1 min to eliminate the previous sample and to ensure that fresh sample was collected. At the same time, the mixed chamber and tubing were washed continuously using filtered PBS in order to avoid crossing contamination. The next step was to feed the mixed chamber with 500 µl of PI diluted in filtered PBS at a concentration of 4 mg/L and adjust the given dilution sequence. For the dilution 1:2, only 500 µl of fresh sample was delivered to the mixed chamber. For the next dilution sequences, the sample was diluted in mixed PBS and PI. The dilution sequences set in this study are ranging from 32 to 8192 by a factor of 2. Then, the mixed sample with PI was then incubated at room temperature for 1 min (for E coli) and 3 min (for pseudomonas) strains. Finally, the sample is automatically transferred to C6 FC (BD Accuri C6, BD Biosciences) and is analyzed at a medium flow rate (33 μl/ min) with a threshold FSC-H set at 16000 and 80000 for *P. putida* and *E. coli* strain respectively. The analysis ended after collecting at least 20,000 events. For each sample, the data are then stored and exported in .xlsx and .fsc formats. All the data related to the different parameters (mean, median, CV) are available to be displayed in real time during the cultivation.

During sample acquisition, the concentration of the number of events per µl was maintained in the range 500–1500 via a script Matlab program to further avoid doublet detection. Indeed, if the number of cells per µl corresponding to the current sample was in the range, the actual dilution rate will be maintained for the next sample. Otherwise, the dilution rate will be increased or reduced by a factor of 2 if the number of cells per µl was above the upper threshold or under the lower threshold respectively.

The behavior of the feedback control loop was controlled by a script Matlab program which is running automatically after the generation of a new data. Thus, cells were gated based on forward scatter (FSC) and FL3 channel (relative to red fluorescence detection) and expressed as a percentage. The lower threshold on FL3 channel has been fixed as the background of fluorescence while the upper threshold is defined as the minimum detectable signal for dead cells. These are the limits of so called dynamic range (DR).

The dynamic cell fraction was modulated by digital control system composed of a peristaltic pump (Watson Marlow, 101 UR) which automatically activated to feed the bioreactor with a pulse of 0.3 g of glucose to maintain the percentage of cells computed in DR at a specified set point (10%).

For every sample, the data are exported as .fsc file and are processed using Matlab software (fca_readfcs.m function by L. Balkay, University of Debrecen, Hungary, available on MatLab central file sharing). In short, the gate was designed as described above to compute the percentage of cells in the dynamic range.

For the preparation of non-viable cells to be used as a positive control for PI staining and flow cytometry gating, 1 ml of cell suspension was heated at 80°C for 1 hour. The cells were then washed and re-suspended in filtered PBS. Then, 5 µl of propidium iodide (1mg/ml) was added to the cell suspension and then incubated for 10 min at room temperature. The red fluorescence signal was measured using the C6 FCM using the parameters described above.

### Poisson process simulation

In order to determine if phenotypic diversification process follows a Poisson processes, several simulations have been carried out. Control of population diversification in segregostat has been made based on glucose pulsing, period with glucose pulses corresponding to active phenotypic diversification. The rate of glucose pulse addition λ (h^-1^) has thus been used for running stochastic simulations based on a Poisson process. All calculations have been made based on Matlab (R2014b) and R.

### Case studies: membrane permeabilization dynamics in *E. coli* and *P. putida* upon nutrient limitation

One of the critical steps for studying the outcome of phenotypic diversification relies on the identification of relevant single cell proxies. In this work, we have identified propidium iodide (PI) as an effective biomarker for cells switching to adaptation in nutrient limitation (Figure 2). PI will be used in order to keep track of the progressive permeabilization of the OM upon nutrient limitation. Indeed, we have observed that, during switch from exponential phase to stationary phase in flasks culture experiments, Gram-negative bacteria undergo OM permeabilization, leading to an intermediate PI-stained fraction of cells (Figure 2B for *E. coli* and Figure 2C for *P. putida*).

Since this particular phenotype seems to be triggered by nutrient limitation, chemostat experiments at low dilution rate, i.e. D = 0.1 h^-1^, have been carried out. Indeed, it has been previously observed that cultivating *E. coli* at this dilution rate triggers adaptation to nutrient limitation, notably through a complete remodeling of the porins at the level of the OM [29]. It has been shown previously that this dilution rate is in the range of growth rate for which active porin remodeling takes place [30]. On this basis, chemostat experiments have been performed both for *E. coli* and *P. putida* with on-line flow cytometry profiling (Figure 3). Similar trends have been observed for *E. coli* and *P. putida*. In the first phase with increasing OM permeabilized subpopulation, followed by a second phase where this subpopulation decreases. Other interesting observation for each species, is a continuous evolution at the level of their subpopulation ratio, whereas chemostat is typically used for “stabilizing” microbial population [REF]. In front of the results, *E. coli* exhibits a lower phenotypic diversification rate than *P. putida*. Indeed, if we compute the rate of diversification from the first phase of chemostat where OM permeabilized subpopulation increases, the value is 0.044 h^-1^ for *E. coli* against 0.085 h^-1^ for *P. putida*. Since chemostat is the standard cultivation tool that is in use in a lot of research laboratories, the fact that it is not possible to maintain cell population heterogeneity at a given level in this device is a very interesting observation, suggesting that data obtained based on chemostat cultivation might need to be reanalyzed in light of the presence of phenotypically different subpopulations.

**Figure 3:**
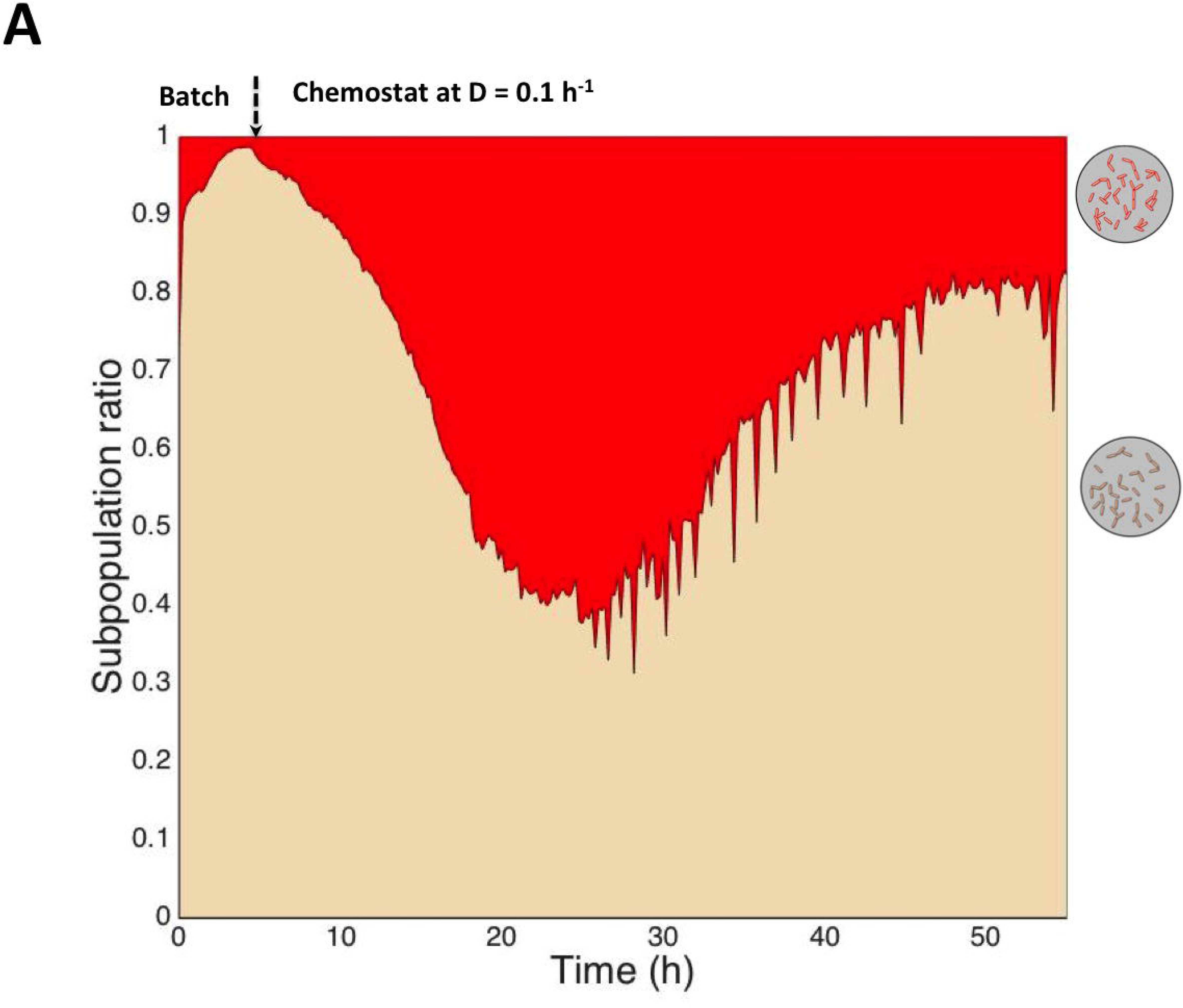
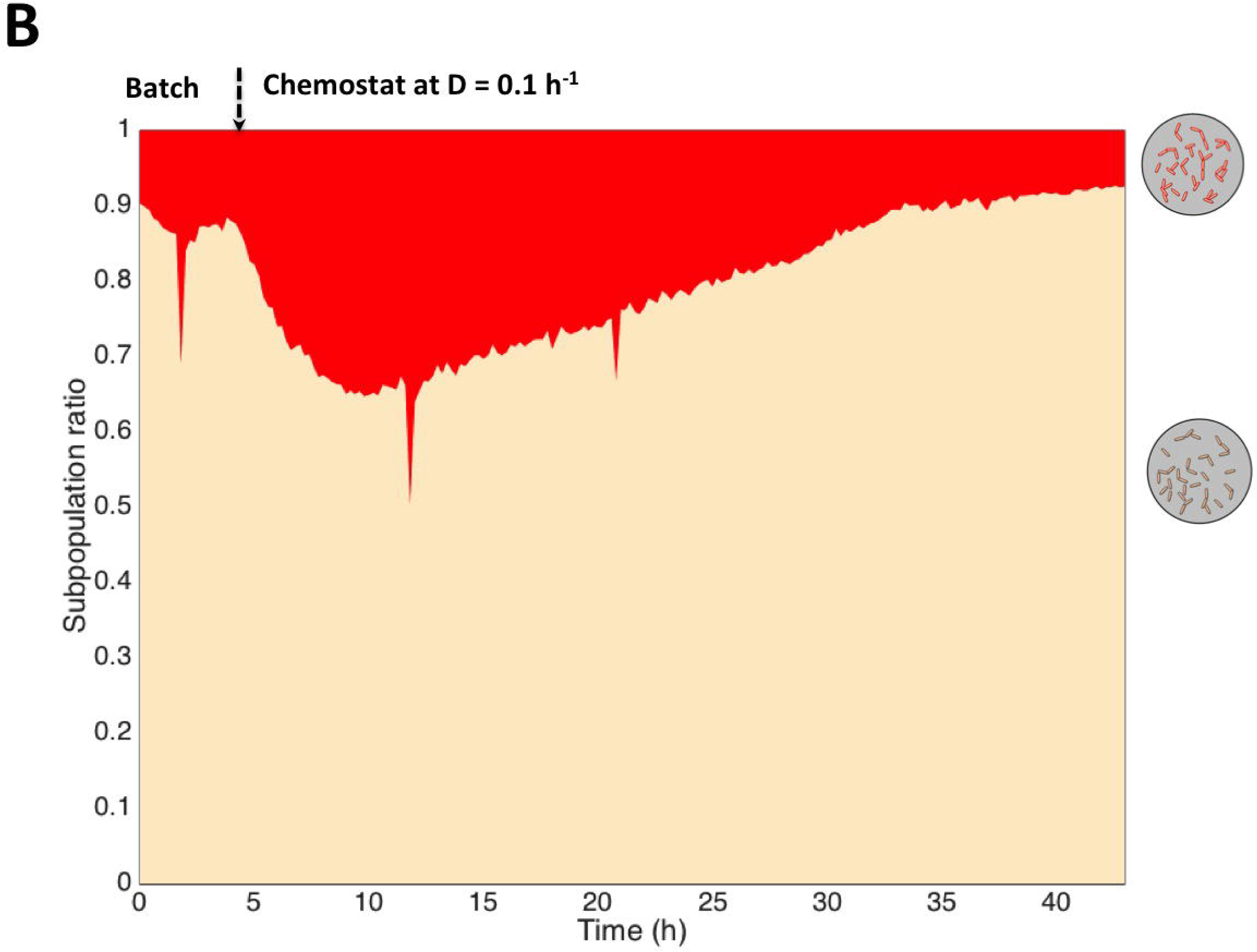
dynamics of phenotypic diversification in chemostat for A: *E. coli*;B: *P. putida.*

Based on these data, we designed an alternative cultivation device for maintaining cell population heterogeneity (or cell subpopulation ratio) at a constant level with time. This device, which we called Segregostat (see Figure 1 for a description of the set-up), is also running in continuous mode, but is under the control of the degree of diversification of the microbial population, i.e. in our case the ratio between cells in non-permeabilized and permeabilized subpopulation. Continuous monitoring of cell population is ensured by on-line flow cytometry like in the chemostat experiments. However, in the case of the segregostat, the system is continuously fed with base medium without any carbon source. The carbon source, i.e. glucose, is injected by pulse at given time interval based on the outcome of flow cytometry analysis. Since the switch of cells to the permeabilized state is triggered by carbon limitation, glucose pulse can be used as an efficient actuator for limiting this phenotypic switch. During the experiments, glucose pulse is triggered automatically when the amount of cells in the permeabilized subpopulation exceed 10% of the total amount of cells (displayed as a subpopulation ratio of 0.1 on figure 4). This threshold has been considered here as being a significant proxies for the induction of the diversification process.

**Figure 4:**
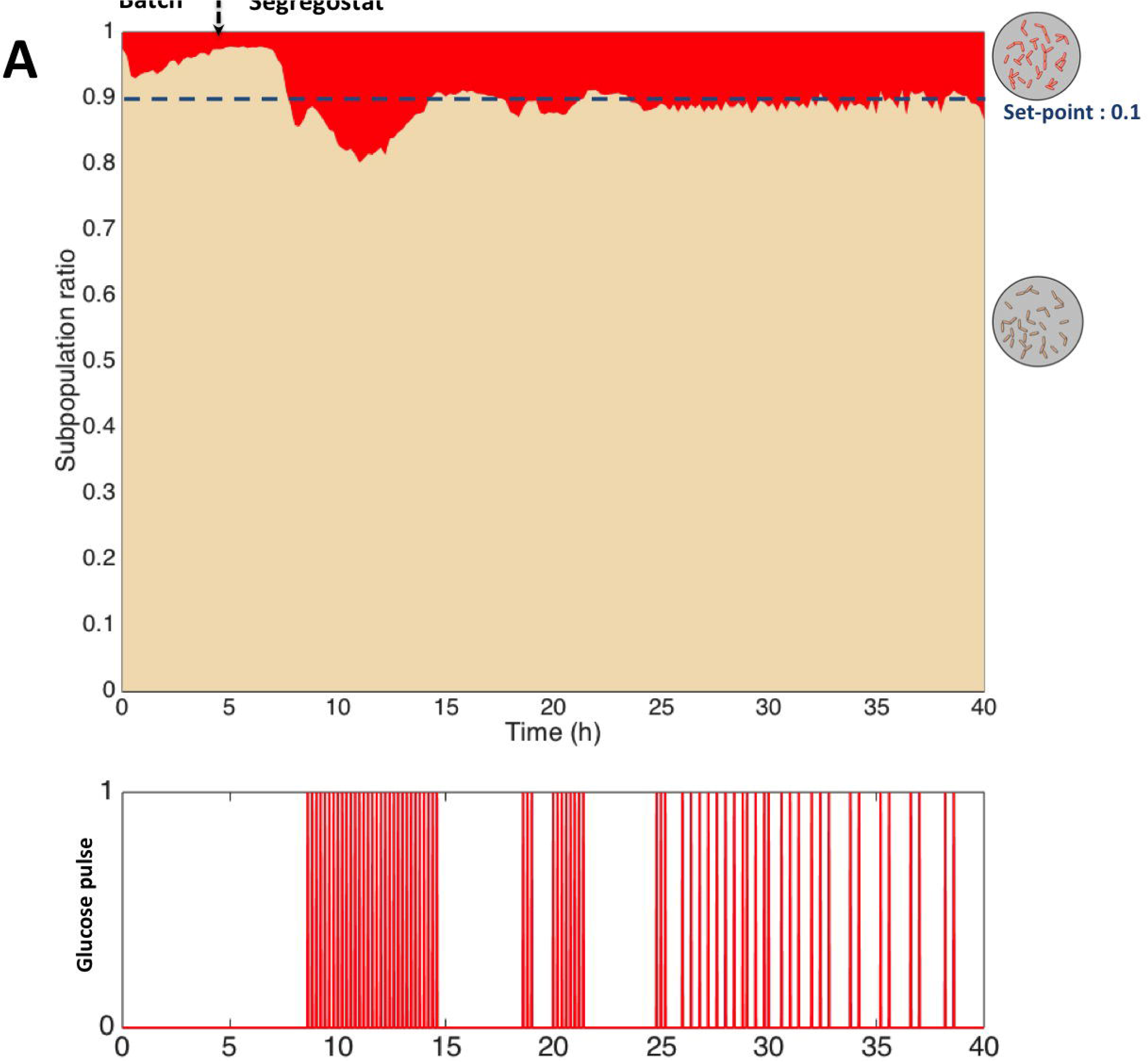
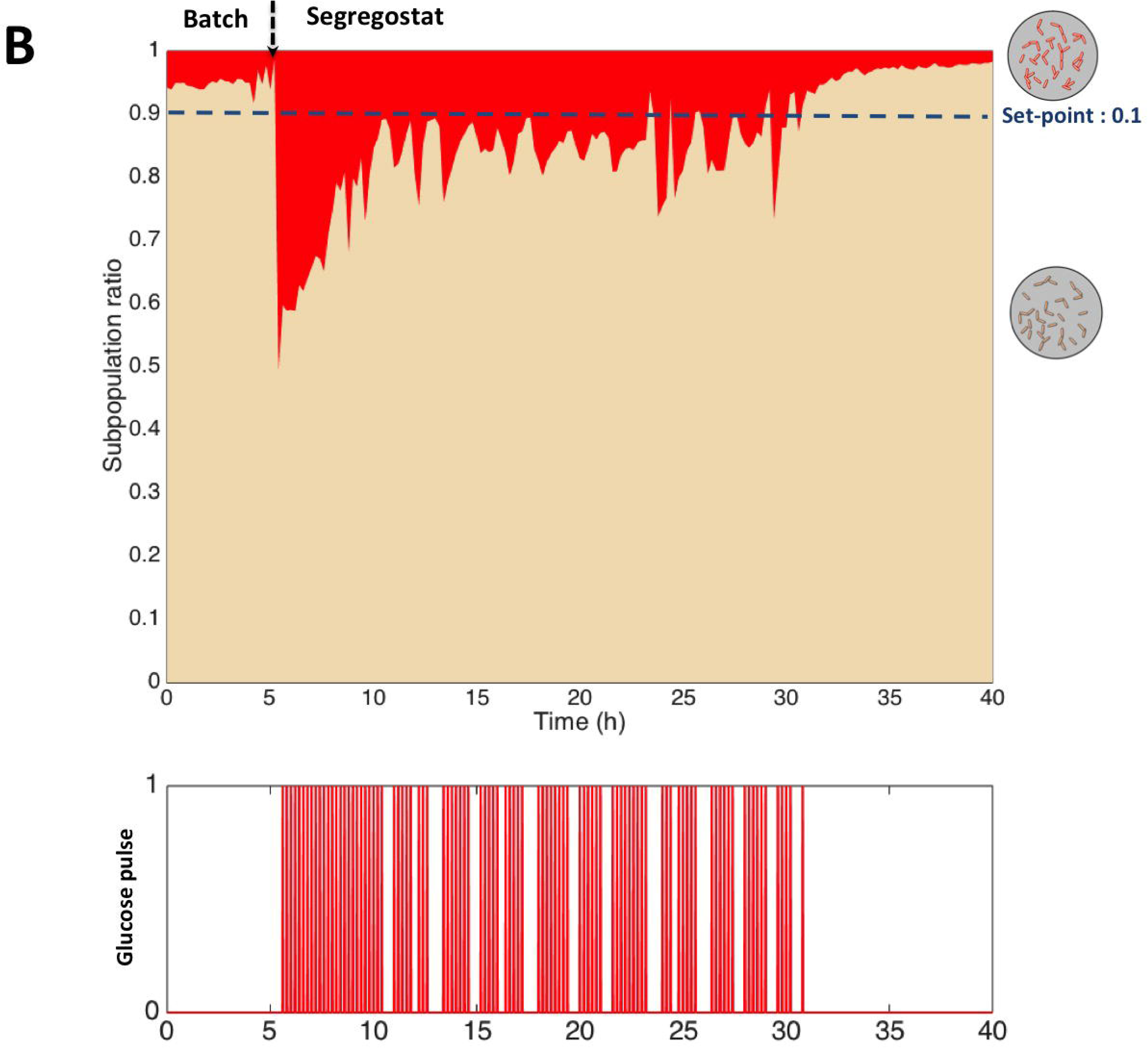
dynamics and control of phenotypic diversification in segregostat for A: *E. coli*; B: *P. putida.* Glucose pulses have been made based on a population diversification ratio of 0.1.

The *E. coli* and *P. putida* profiles present clear similitudes, but also some differences. First, upon the entry in the segregostat mode, both Gram-negative bacteria exhibit a strong phenotypic diversification traduced by a series of glucose pulses (Figure 4; ∼5h-10h]. However, this process seems to be faster for *P. putida* than for *E. coli*. Indeed, a nice feature of the segregostat is that the diversification periods can be easily tracked based on the glucose pulse profile. Indeed, these Gram-negative bacteria tend to induce OM permeabilization upon nutrient limitation, this process being controlled by pulsing glucose during the culture. Then, a period with successive glucose pulses corresponds to a period of intense phenotypic diversification. The process is faster for *P. putida*, with an average 3.9 pulses/h, by comparison with *E. coli* for which the average is 2.28 pulses/h. these observations are in good accordance with the results gained through chemostat experiments that have shown that the diversification rate is approximately twice as much for *P. putida* than for *E. coli*.

At this level, it should be interesting to determine whether the pulses addition follow a stochastic trend. For this purpose, stochastic simulations based on Poisson processes have been performed based on the transition rates (λ, in h^-1^) experimentally determined through the segregostat experiments (i.e., 2.28 h^-1^ for *E. coli* and 3.9 h^-1^ for *P. putida*).

Poisson processes have been simulated based on the average number of pulses per hours and the total pulses injected over the segregostat cultures for the two Gram-negative models (Figure 5A). Interestingly, it can be observed that the mean time computed from the distribution, i.e. 24.9 h, match very well with the experimental time which is roughly around 25 h.

**Figure 5:**
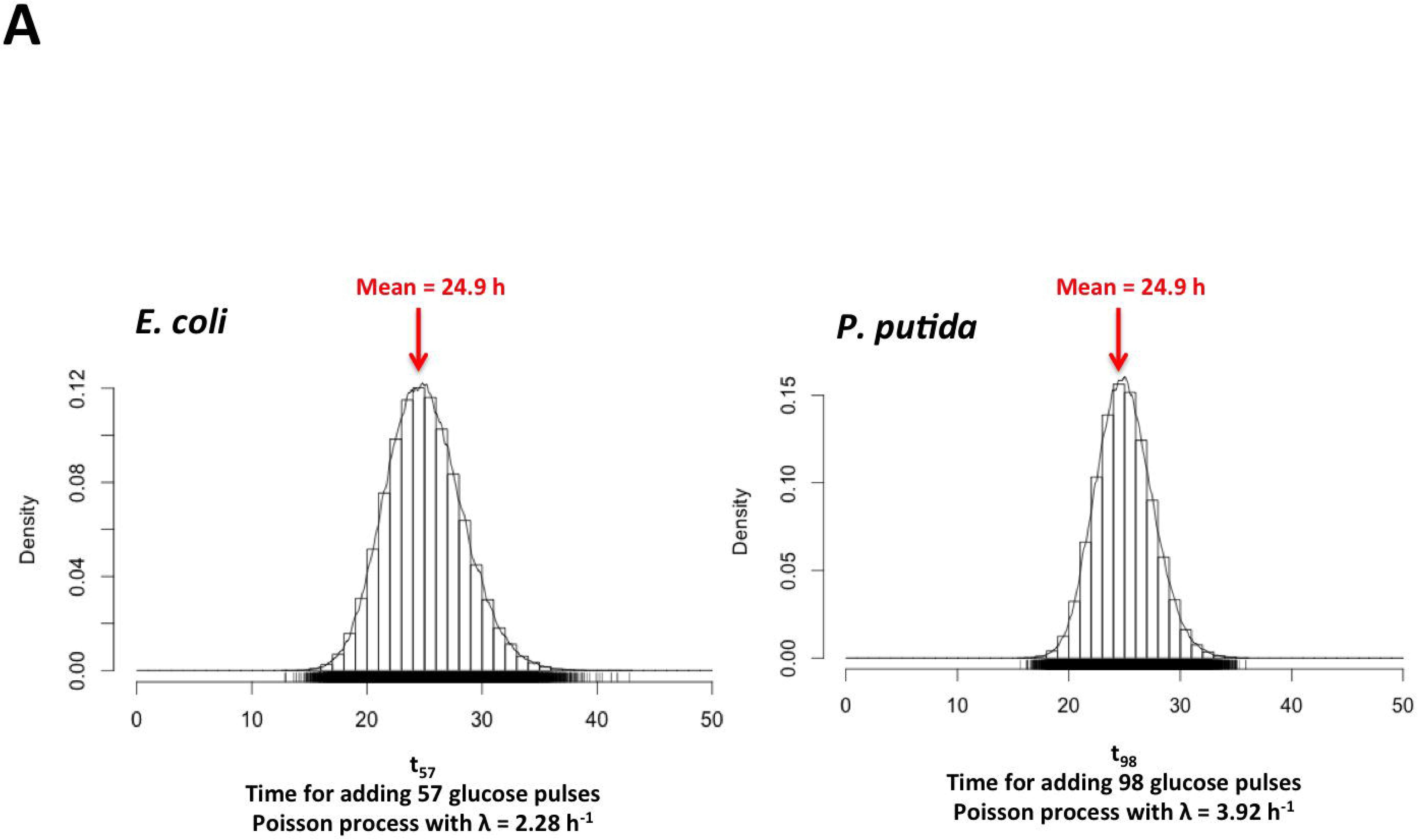
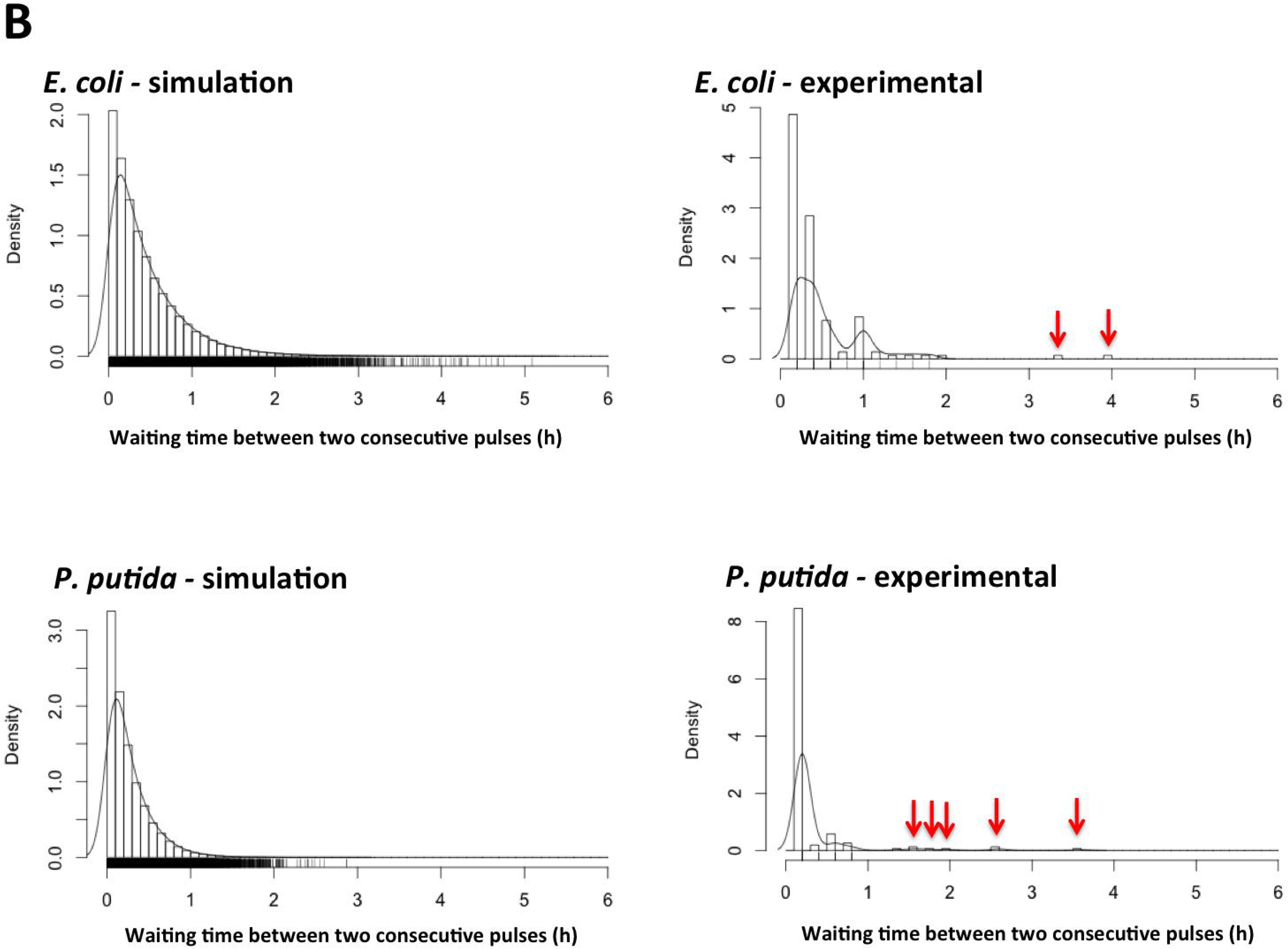
analysis of glucose pulses addition profiles in segregostat A Total time needed for adding 57 and 98 glucose pulses respectively according to a theoretical Poission process (the number of pulses, as well as the rate λ of the Poisson process have been determined from the experimental segregostat data). B distribution of waiting time between two consecutive glucose pulses according to a theoretical Poisson law (simulation) and from the segregostat data (experimental)

However, when looking at the glucose pulse profiles displayed in figure 4, it can be observed that we are diverging from a classical Poisson process. It is particularly obvious in the case of *P. putida* where bursts in glucose pulses are observed, i.e. period with successive pulses followed by period with no pulses. In order to highlight this phenomenon, we have computed the waiting time distribution, i.e. in our case time between two successive glucose pulses, from stochastic simulations involving Poisson processes with the transition rate determined experimentally (i.e., again 2.28 h^-1^ for *E. coli* and 3.9 h^-1^ for *P. putida*) and made a comparison with the distribution that have been acquired experimentally (Figure 5B). There are two interesting features that need explanation. The first feature relies on the fact that, in the experimental distribution, a lot of low waiting time values are non represented whereas some others are overrepresented. This is due to the fact that in the segregostat set-up, glucose pulses are added in discrete time intervals (glucose pulse is potentially injected every 12 minutes depending on the degree of diversification of the microbial population), whereas stochastic simulations are carried out in continuous-time. The second, and most interesting value, is that some high waiting time values, not predicted by stochastic simulations, can be noticed in both cases, but the effect is more obvious for *P. putida* (these values has been pointed out by red arrows for the experimental distribution displayed in figure 5). These values correspond to the arrests in glucose pulses profiles displayed in figure 4 and are typical of a bursting process, i.e. process exhibiting period of intense diversification, followed by periods with no activities. Possible explanations about the source of such burst behavior will be discussed in the next section.

## Discussion

Phenotypic diversification is due to stochasticity in gene expression, arising from fluctuations in transcription and translation despite constant environmental conditions. These mechanisms have been the focus of intensive researches during the last decade and have led to a coherent mathematical and experimental framework of molecular stochasticity in prokaryotic and eukaryotic systems [31][3]. This framework has been notably used in order to decipher the impact of regulatory network structure on the propagation [32][8] and control of phenotypic diversification [31][33][34], as well as on the possible functionality of such diversification [3][35][10]. However, most of these researches have been conducted at low spatio-temporal resolution, i.e. either on a limited numbers of cells, or focused on given time point. In this work, we have developed a tool, called segregostat, allowing to expand the methodology to a very high amount of cells with a high temporal resolution. Indeed, the segregostat allowed generating high-quality subpopulation data. Such data are rarely available at such population density and should help paving the way for further characterization of microbial cell population diversification dynamics. One characteristic feature of the version of the segregostat that has been shown in this work is that population control is made based on PFM. The resulting glucose pulse profile can be then used for identifying the period of time for which intense phenotypic diversification occurs. Interestingly, it has been shown that the time between two consecutive glucose pulses was not following a simple Poisson process, but was rather occurring in a burst fashion. Transcriptional and translational bursting are known to occur at the single cell level and drive intrinsic noise leading to phenotypic diversification [3][31]. However, this process has never been described for a whole microbial population and an outstanding question would be to understand how such typical single cell dynamics can be transposed to the behavior of whole microbial population. One hypothesis is that glucose pulsing resulting from PFM results in synchronization in microbial subpopulations dynamics during cultivation in chemostat. Indeed, since control is made based on glucose pulsing, microbial population is exposed to cycles of nutrient excess and limitation, based on a given diversification state. This hypothesis needs of course confirmation and extra work is still required in order to fully understand the molecular mechanisms behind the appearance of OM-permeabilized subpopulation.

We have shown that *E. coli* and *P. putida* exhibit totally different dynamics. For *E. coli*, a simple PMC strategy can be used for controlling phenotypic diversification rate and maintaining microbial population in a given degree of heterogeneity with time. For *P. putida*, PMC leads to oscillations around the subpopulation ratio that has been used as the set-point for the pulse addition, and eventually lead to a lost of control at the end of the cultivation. It seems thus that, in this case, an advanced controlled strategy based on predictive model, will be required. From a mechanistic point of view, glucose uptake mechanisms and catabolism reactions between *E. coli* and *P. putida* are very distinct. These differences could in part be responsible for the observed dynamics. In *E. coli* all the glycolytic reactions take place in the cytoplasm. In contrast, *P. putida* carries out some of the reactions leading to glucose catabolism in the periplasm. Indeed, it has been shown that small amounts of gluconate and 2-ketogluconate accumulate in the extracellular space when *P. putida* grows on glucose [26].

From a functional perspective, population heterogeneity in membrane permeabilization state can be implied in fundamental processes, such as stress resistance (bet-hedging), increase of nutrient uptake capacity, or on the opposite extended leakage of compounds implied in cross-feeding and microbial interactions. The methodology presented in this work and relying on the use of segregostat will be an important component of the experimental and numerical workflow needed for addressing these very complex physiological processes.

This work points out the importance of choosing an appropriate single cell proxy for controlling phenotypic diversification and that the resulting on-line profiling of subpopulations can lead to a more thorough understanding of population dynamics.

In conclusion, the segregostat can in future be used for several potential applications raging from: generating population of constant structure over time to be used for extended physiological studies, controlling bioprocesses and “homogenize” bioprocess populations, analyzing and controlling synthetic co-culture processes, and at a global level, expand our knowledge about the dynamics of phenotypic diversification of microbial populations with possible link to the functionality of this diversification process.

## Supporting information

Supporting informations

## Acknowledgement

HS, FD are supported by a Wagralim-Biowin grant (SingleCells project n° 7273). TMN is supported by a grant provided by the Vietnamese government (VIED).

