## Supporting informations for "Segregostat: A novel concept to control phenotypic diversification dynamics on the example of Gram-negative bacteria"

As stated in the main document, *E. coli* [JW2203-1](http://cgsc.biology.yale.edu/Strain.php?ID=108062) Δ*ompC* have been selected based on a prescreening test since it was able to display a higher diversification ratio by comparison with wild type and other porin mutants (i.e. Δ*ompF* and Δ*lamB*). All these strains (table S1) have been stained with PI and analyzed by FC (Figure S1).

### Cultures were performed in Bioscreen C microplate reader at 37°C on a defined mineral salt medium containing (in g/L): K_2_HPO_4_ 14.6, NaH_2_PO_4_.2H_2_O 3.6, Na_2_SO_4_ 2, (NH_4_)_2_SO_4_ 2.47, NH_4_Cl 0.5, (NH_4_)2-H-citrate 1, glucose 5, thiamine 0.01, kanamycin 0.1. Thiamine and kanamycin were sterilised by filtration (0.2μm). The medium is supplemented with 3 mL/L trace solution, 3 mL/L FeCl_3_.6H_2_O solution (16.7 g/L), 3 mL/L EDTA solution (20.1 g/L) and 2 ml/L MgSO_4_ solution (120 g/L). The trace solution contains (in g/L): CoCl_2_.H_2_O 0.74, ZnSO_4_.7H_2_O 0.18, MnSO_4_.H_2_O 0.1, CuSO_4_.5H_2_O 0.1 and CoSO_4_.7H_2_O 0.21.

**Table S1:** list of strains used in this work

| **Name** | **Genotype** | CGSC# |
| --- | --- | --- |
| [BW25113](http://cgsc.biology.yale.edu/Strain.php?ID=64667) WT | F-, *Δ(araD-araB)567*, *ΔlacZ4787*(::rrnB-3), *λ^-^*, *rph-1*, *Δ(rhaD-rhaB)568*, *hsdR514* | 7636 |
| [JW3996-1](http://cgsc.biology.yale.edu/Strain.php?ID=109158) *ΔlamB* | F-, *Δ(araD-araB)567*, *ΔlacZ4787*(::rrnB-3), *λ^-^*, *rph-1*, *Δ(rhaD-rhaB)568*, *ΔlamB732::kan*, *hsdR514* | 10877 |
| [JW0912-1](http://cgsc.biology.yale.edu/Strain.php?ID=107206) *ΔompF* | F-, *Δ(araD-araB)567*, *ΔlacZ4787*(::rrnB-3), *λ^-^*, *ΔompF746::kan*, *rph-1*, *Δ(rhaD-rhaB)568*, *hsdR514* | 8925 |
| [JW2203-1](http://cgsc.biology.yale.edu/Strain.php?ID=108062) *ΔompC* | F-, *Δ(araD-araB)567*, *ΔlacZ4787*(::rrnB-3), *λ^-^*, *ΔompC768::kan*, *rph-1*, *Δ(rhaD-rhaB)568*, *hsdR514* | 9781 |

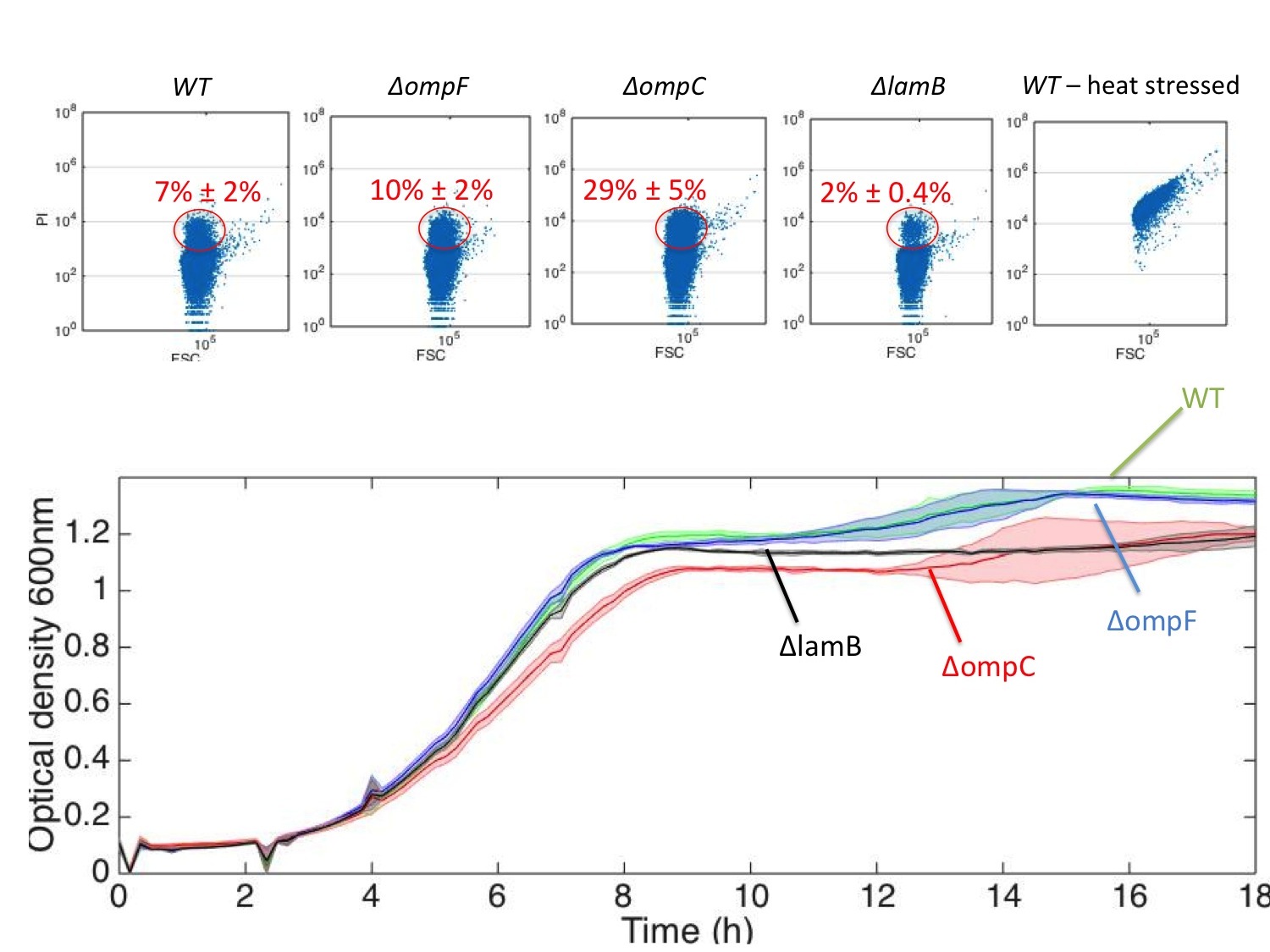

**Figure S1** : (TOP) permeabilization of OM of E. coli leads to the accumulation of PI in the perisplasmic space. Flow cytometry analyses (x-axis : FSC, i.e. forward scatter proportional to cell size ; y-axis : PI level, i.e. fluorescence related to PI accumulation in cells) of wild type (WT), deletion mutants and wild type exposed to heat stress (60°C for 30 minutes). (BOTTOM) Evolution of optical density in Bioscreen C microplate cultivation device (cultures have been made in triplicates, mean and standard deviation are indicated). Samples have been taken after 18h of cultivation in Bioscreen C, stained with PI and analyzed by FC.
